# Genome-wide comparisons and extrapolations of AU-rich elements in Plants with *Homo sapiens*

**DOI:** 10.1101/2020.04.28.066589

**Authors:** Deepshikha Satish, Anil Panwar

## Abstract

The Adenylate-Uridylate Rich elements (AREs) are adenine and uracil abundant sequences, established in ephemeral mRNAs, principally present in the 3’ untranslated region (3’ UTR). AREs are widely accepted as a cause of high turnover of the mRNAs containing them (Bakheet, 2001; Barreau, 2005; Shaw and Kamen, 1986). The mammalian ARE-mRNAs primarily translate into nuclear transcription factors, oncoproteins, cytokines, and G-protein-coupled receptors which indicates their indispensable role in the regulation of gene transcription during cell growth and differentiation, and the immune response (Chen and Shyu, 1995; Wilson et al., 1999). The present study is an attempt to comprehensively analyze the eloquent presence of AU rich elements in genome, transcriptome and 3’UTR of three important plant species (*Arabidopsis thaliana, Oryza sativa and Zea mays*) and compare the statistics of putative ARE motifs in plants with *H.sapiens*. Statistical analysis of genome-wide putative AU-rich elements revealed the explicit presence of ARE motifs in plants. This is the first study that analyses the presence of Adenylate-Uridylate Rich elements (AREs) in three different plants which can be further validated through experiments.

## Introduction

The management and control of mRNA stability is of utmost essentiality for adjusting mRNA abundance and for restricting mRNA translation to prevent detrimental effects caused by overexpression of proteins (Caput et al., 1986). Adenylate-Uridylate Rich elements (AREs) are nucleotide sequences exuberant in adenine and uracil, present within the 3’ untranslated region (UTR) of ephemeral mRNAs and are liable for high turnover of the mRNA containing them (Bakheet, 2001; Barreau, 2005; Shaw and Kamen, 1986). ARE containing mRNAs encode proteins that regulate either cell growth or the impedance of an organism to external factors like micro-organisms, inflammatory stimuli and environmental factors. Such genes require meticulous control of their temporal and spatial expression patterns which is accomplished, in addition to transcriptional control, by a regulation of the mRNA translation and therefore the stability of the mRNA a. In mammalian cells, ARE-mRNA principally codes for nuclear transcription factors, oncoproteins, cytokines, and G-protein-coupled receptors which specify their indispensable role in the modulation of immune reaction, gene transcription during cell growth and differentiation (Chen and Shyu, 1995; Wilson et al., 1999).

So far there is no universal consensus for ARE motifs but the scientific experiments illuminate a list of motifs which are predicted to promote mRNA degradation (Table1). Significant characteristics of ARE elements include the occurrence of several AUUUA pentamers (short distance from each other is preferred) and theoretically the pentamers are implanted within a broader U-rich or AU-rich context. Further, the minimal functional ARE sequence is the nonamer UUAUUUA(U/A)(U/A) in recurrent instances. The three broad classes of AREs according to Chen and co-workers can be recognized based on the subsequent sequence characteristics: class I—one or more AUUUA pentamers within a U-rich sequence; class II— tandem repeats of AUUUA; and class III—U-rich sequences lacking AUUUA motifs (Chen and Shyu, 1995; Wilusz et al., 2001). On the other hand, according to Bakheet’s method, AREs are classified into five groups based on the number of continuous repeats of the core sequence of AUUUA. Groups 1, 2, 3 and 4 possess 5, 4, 3, and 2 continuous repeats of AUUUA, respectively, whereas group 5 AREs possess only one core pattern in the transcript (Bakheet, 2001).

**Table 1:**
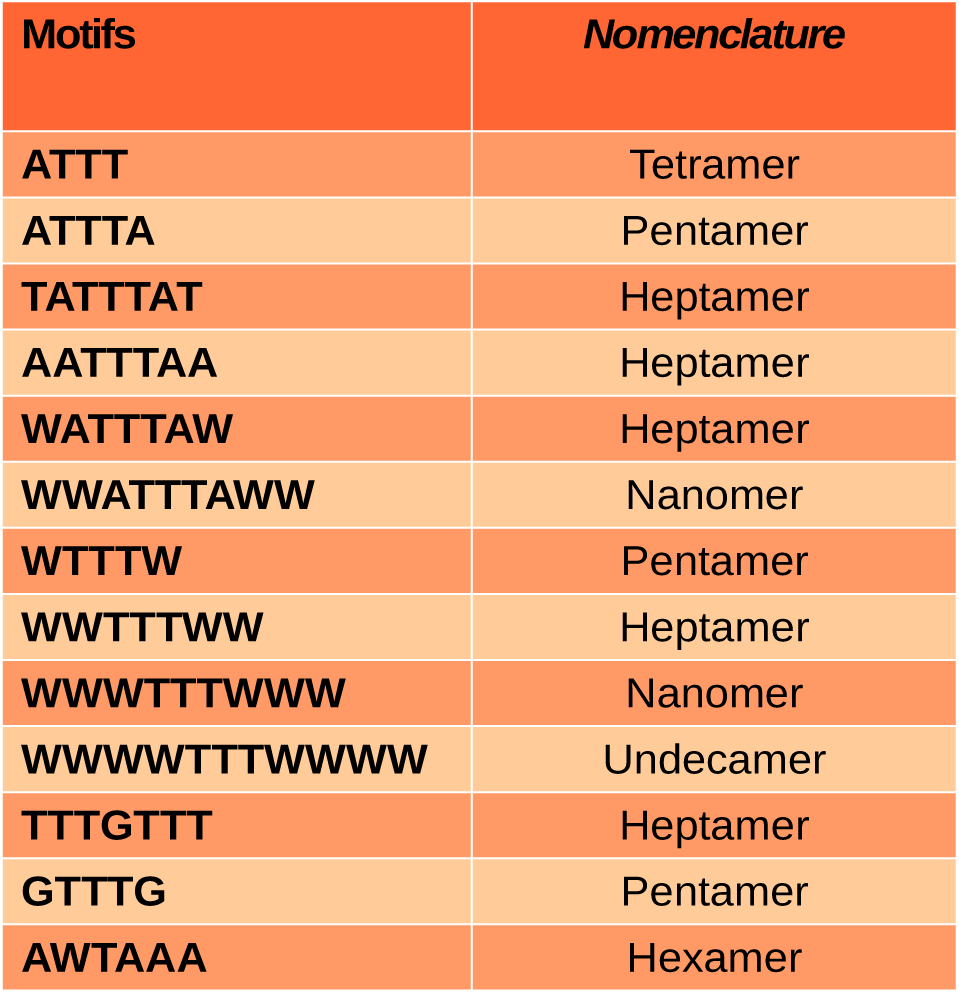
Significant AU rich elements motifs.

These AREs belonging to different classes becomes functional only after interacting with AU-binding proteins (AUBPs) which recruit enzymatic machinery responsible for mRNA degradation in a process described as ARE-mediated decay (Barreau, 2005). Several different proteins (e.g. HuA, HuB, HuC, HuD, HuR) bind to these elements and stabilise the mRNA while others (AUF1, TTP, BRF1, TIA-1, TIAR, and KSRP) destabilize the mRNA, miRNAs may also bind to some of them. These proteins act as adapters that recognize the ARE as a signal and couple the bound mRNAs to the general mRNA destabilization/stabilization machinery. The activity of the AU-element binding proteins is regulated, usually by phosphorylation but also by subcellular localization. In the cytoplasm, AUBPs (like TTP) bind to ARE (AUUUA) in the mRNA. The binding of AUBPs recruits either Dcp, which promotes decapping of mRNA, or deadenylase, which removes the poly (A) tail. After exposure of terminal mRNA, exonucleases act to degrade the mRNA in either the 5′–3′ direction using Xrn1 or in the 3′–5′ direction using the exosome. The preferable mode of action is mediated by rapid deadenylation (poly(A) tail decay) and subsequent degradation of the mRNAs (Figure 1)

**Figure 1:**
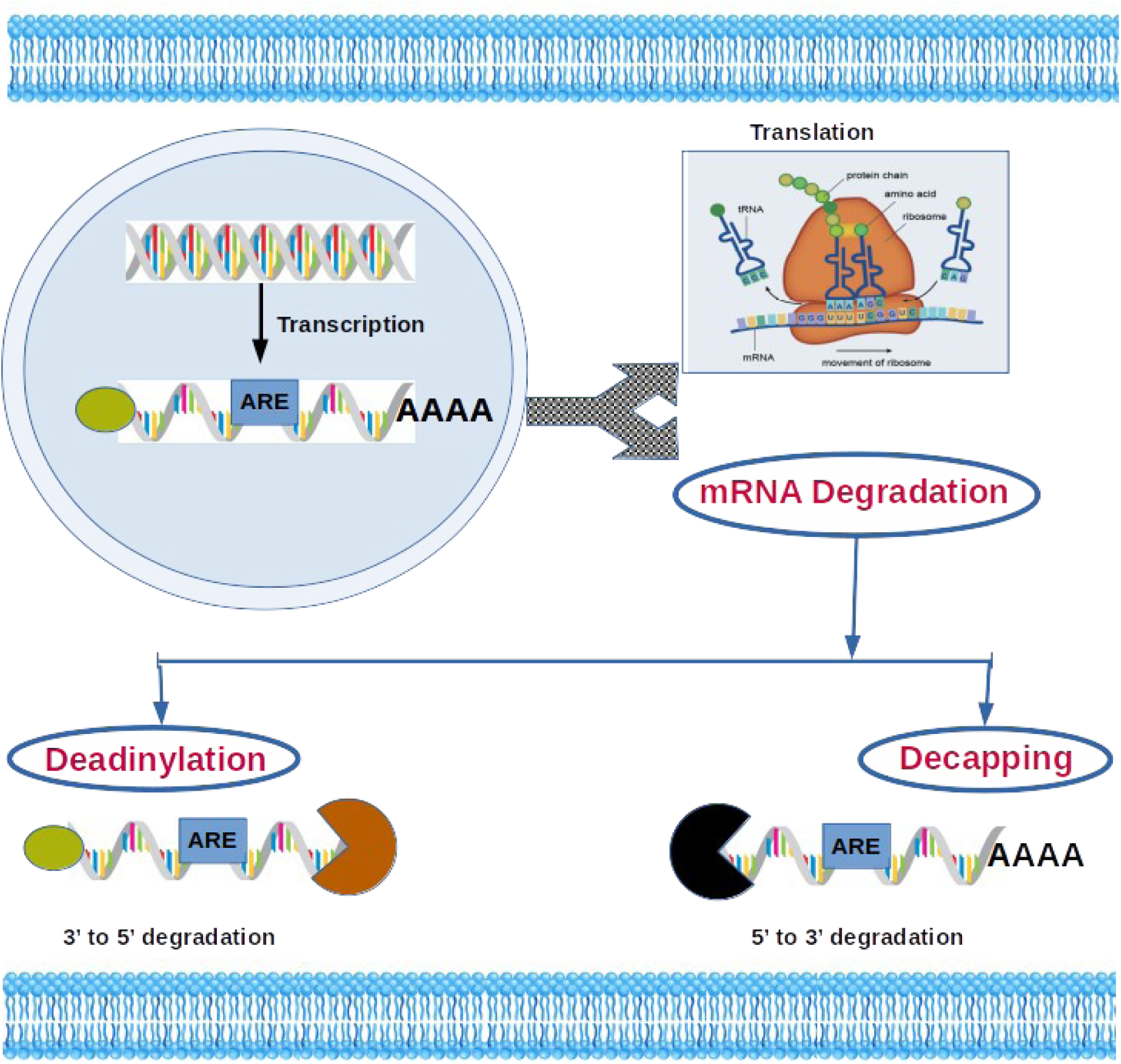
ARE mediated decay of mRNA.

A significant study illustrate that the consistency of AREs is high among the most unstable mRNAs (half-life <2 h), while its existence decreases among mRNAs with enhancing half-life (Lam et al., 2001). Generally, AREs cause mRNAs to be degraded rapidly and thereby suppress gene expression at the post-transcriptional level. Thus, stabilization of the ARE-mRNAs in some cases can cause a prolonged response that may subsequently lead to a diseased state. Problems with mRNA stability have been identified in viral genomes, cancer cells, and various diseases. Accumlating evidences indicate that many of these problems arise because of faulty ARE function. For instance, the *c-fos* gene produces a transcription factor that is activated in several cancers and it lacks the ARE elements. *c-myc* gene, also responsible for producing transcription factors found in several cancers, has also been reported to lack the ARE elements. The *Cox-2* gene catalyses the production of prostaglandins—it overexpresses in several cancers and is stabilized by the binding of CUGBP2 RNA-binding protein to ARE. The association between turnover of defective mRNA and diseases underscores the significance of a deep understanding of the ARE-mediated decay mechanism (Eberhardt et al., 2007; Espel, 2005; Khabar, 2005). Yet, a compilation of mRNAs and their AREs along with half-life is still missing in available databases.

AU rich elements are not only one of the most prominent indicator for RNA stability in the mammalian cells (Chen and Shyu, 1995) but, ARE-bearing mRNAs are wide-spread in species ranging from *Homo sapiens* to the *Saccharomyces cerevisiae* (Duttagupta et al., 2003). This widespread presence of AU rich elements and their crucial roles inspire us to extrapolate their cross-kingdom roles. Sporadic reports indicating the presence and role of AREs in plants kingdom are already available but no genome-wide study of ARE in multiple plants species is available yet. This scientific void of knowledge motivates us to predict the genome-wide presence of different ARE motifs in three important plant species at genome, transcriptome and 3’ UTR level and compare them with human AU rich elements statistics. To understand the significance of AREs, computational analysis was carried out to identify the prevalence of AREs in *Arabidopsis thaliana, Oryza sativa* and *Zea mays* mRNAs and compared with *Homo sapiens*. The present study might open a new avenue to further establish the relationship between ARE types, positions and the transcript half-life in the plant kingdom.

## Material and methods

Genomic Data and Transcriptomic data for the present study were retrieved from the NCBI server (https://www.ncbi.nlm.nih.gov/) while 3’ Untranslated Region Data was retrieved from http://utrdb.ba.itb.cnr.it/. Complete genomes of *A. thaliana, O. sativa, Z. mays and H. sapiens* were taken as initiation point. An in-house Perl script was used to deduce a total number of 14 different ARE motifs (Figure 2). Further complete transcriptomic data belonging to the above three plant species and humans were used and the number of different ARE motifs were searched in this data as well. As most of the functional ARE sites reported till now belong to untranslated regions of the genome, hence we tried to estimate the number of different ARE motifs in a comprehensive pool of 3’UTRs belonging to four species under study. All the results obtained were analyzed and materialized using in-house shell scripts.

**Figure 2:**
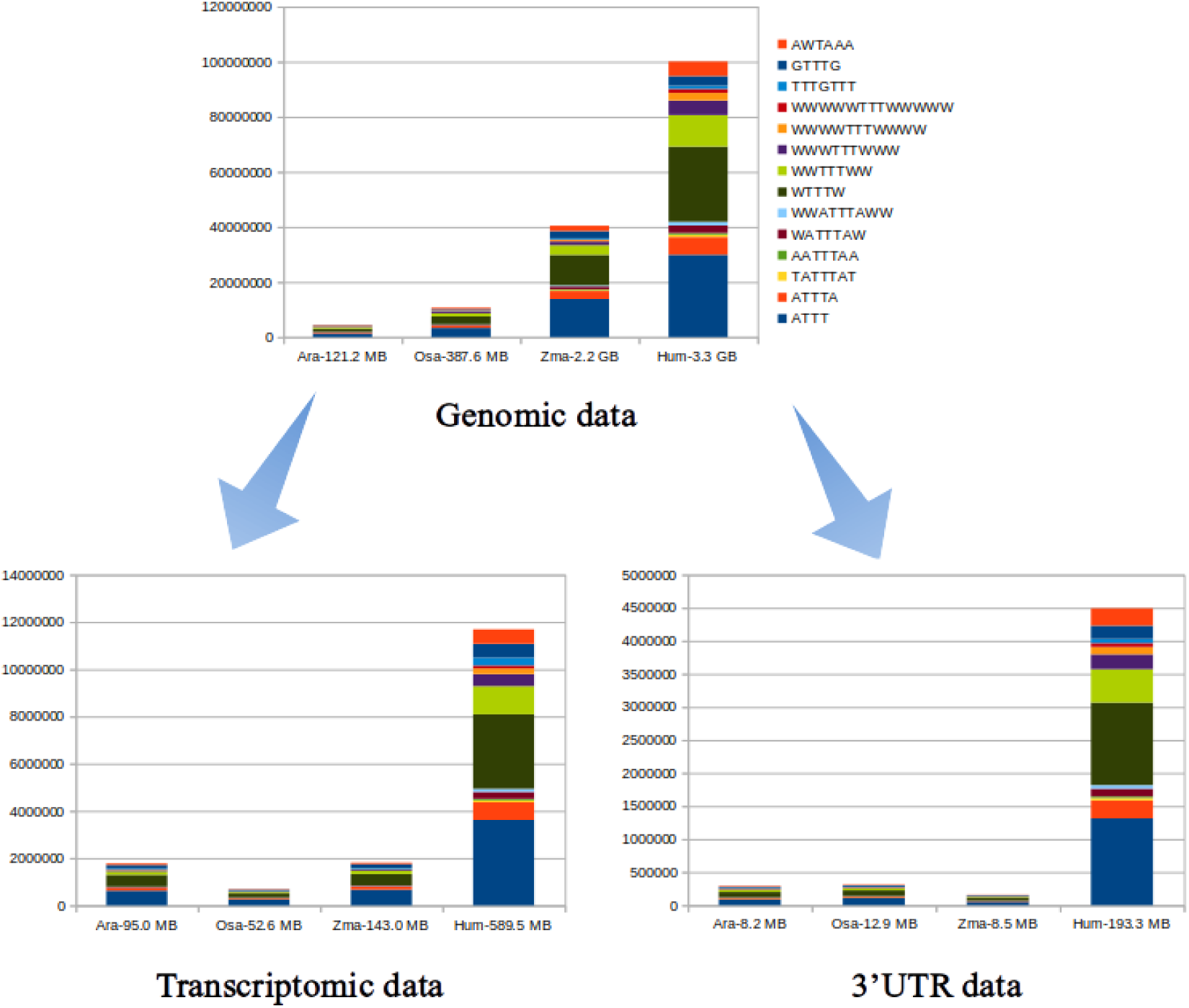
Graphical representation of number of putative ARE motifs predicted in genomes, transcriptomes and 3’UTRs. Ara=Arabidopsis thaliana, Osa= O.sativa, Zma= Z.mays, Hum= H.sapiens. Numbers on the X-axis indicate sizes of respective nucleotide data.

## Results and discussion

Bioinformatics analysis predicted that as many as 7% (≈2000) of the mRNAs in the human transcriptome harbour AREs (Bakheet, 2001). However, very few of these AREs have actually been validated experimentally.

Majority of studies regarding AU rich elements uses animals as the target (Fan et al., 1997; He et al., 2012; Hitti et al., 2016; Spasic et al., 2012) while present study is the first of its kind where the magnitude of putative ARE sites are estimated in plants at the genome-wide scale.

Hitherto, the presence of ARE motifs in 3’UTR only has been taken into cognisance by the various scientific studies but we made attempt to investigate the occurrence of ARE motifs at genomic and transcriptomic colosseum also. This comprehensive analysis made us to draw some interesting conclusions (Figure 2). The ARE motif occurrence in genomic data was anticipated to be directly correlated with the number of nucleotides. But corelation of number of ARE motifs at transcriptomic and 3’UTR scale exhibited some very intriguing inclinations. For instance, the number of different ARE motifs in *A.thaliana* is not the exclusive reflection of the size of *A.thaliana* transcriptome size. Transcriptome size of *A.thaliana* (95.0 MB) is 33.50% smaller than *Z. Mays* (143 MB), regardless of this difference in size, number of ARE motifs found in *A.thaliana* are just 2% lesser than *Z.mays*. Further, this difference seems to be contributed by the overwhelming presence of “WTTTW” and “ATTT” motifs in the transcriptome of *A.thaliana* (Table 2, Table 3, Table 4, Figure 2). Further, this trend got reversed in 3’UTR data. Size of 3’UTR of *A.thaliana* is just 3.65% smaller than *Z.mays* while the number of putative ARE motifs are reported to be 46.1% higher in *A.thaliana* (Table 2, Table 3, Table 4, Figure 2).

**Table 2:** Number of ARE motifs present in genomes of respective plant species and H.sapiens.

| Motifs | *A. thaliana* | *O. sativa* | *Z. mays* | *Homo sapiens* |
| --- | --- | --- | --- | --- |
| ATTT | 1310215 | 3265887 | 13788857 | 29851282 |
| ATTTA | 290586 | 720476 | 2871616 | 6479456 |
| TATTTAT | 35291 | 82237 | 372208 | 843462 |
| AATTTAA | 31894 | 83051 | 237269 | 626299 |
| WATTTAW | 124024 | 309717 | 1069736 | 2707257 |
| WWATTTAWW | 61751 | 146470 | 419257 | 1335749 |
| WTTTW | 1198621 | 2810838 | 11041178 | 27325857 |
| WWTTTWW | 464010 | 1145879 | 3531366 | 11472366 |
| WWWTTTWWW | 214694 | 548704 | 1277612 | 5273269 |
| WWWWTTTWWWWW | 110086 | 271302 | 555394 | 2709335 |
| WWWWWTTTWWWWW | 60330 | 139527 | 247251 | 1527654 |
| TTTGTTT | 64233 | 102180 | 480650 | 1096963 |
| GTTTG | 183720 | 460821 | 2559491 | 3519206 |
| AWTAAA | 227579 | 528285 | 1951519 | 5305867 |

**Table 3:** Number of ARE motifs present in transcriptomes of respective plant species and H.sapiens.

| Motifs | *A. thaliana* | *O. sativa* | *Z. mays* | *Homo sapiens* |
| --- | --- | --- | --- | --- |
| ATTT | 621490 | 251813 | 660984 | 3614954 |
| ATTTA | 105252 | 41320 | 113303 | 742011 |
| TATTTAT | 9971 | 4418 | 12667 | 84071 |
| AATTTAA | 7857 | 3122 | 7172 | 69770 |
| WATTTAW | 32771 | 14145 | 36793 | 292870 |
| WWATTTAWW | 12356 | 5001 | 12591 | 128664 |
| WTTTW | 496558 | 180175 | 476968 | 3153179 |
| WWTTTWW | 139497 | 55828 | 144431 | 1190921 |
| WWWTTTWWW | 50792 | 20978 | 50428 | 509720 |
| WWWWTTTWWWWW | 21144 | 8388 | 18252 | 246544 |
| WWWWWTTTWWWWW | 9773 | 3690 | 7536 | 124387 |
| TTTGTTT | 45076 | 17609 | 43304 | 329354 |
| GTTTG | 149631 | 55170 | 162416 | 584026 |
| AWTAAA | 70352 | 22383 | 61958 | 615665 |

**Table 4:**
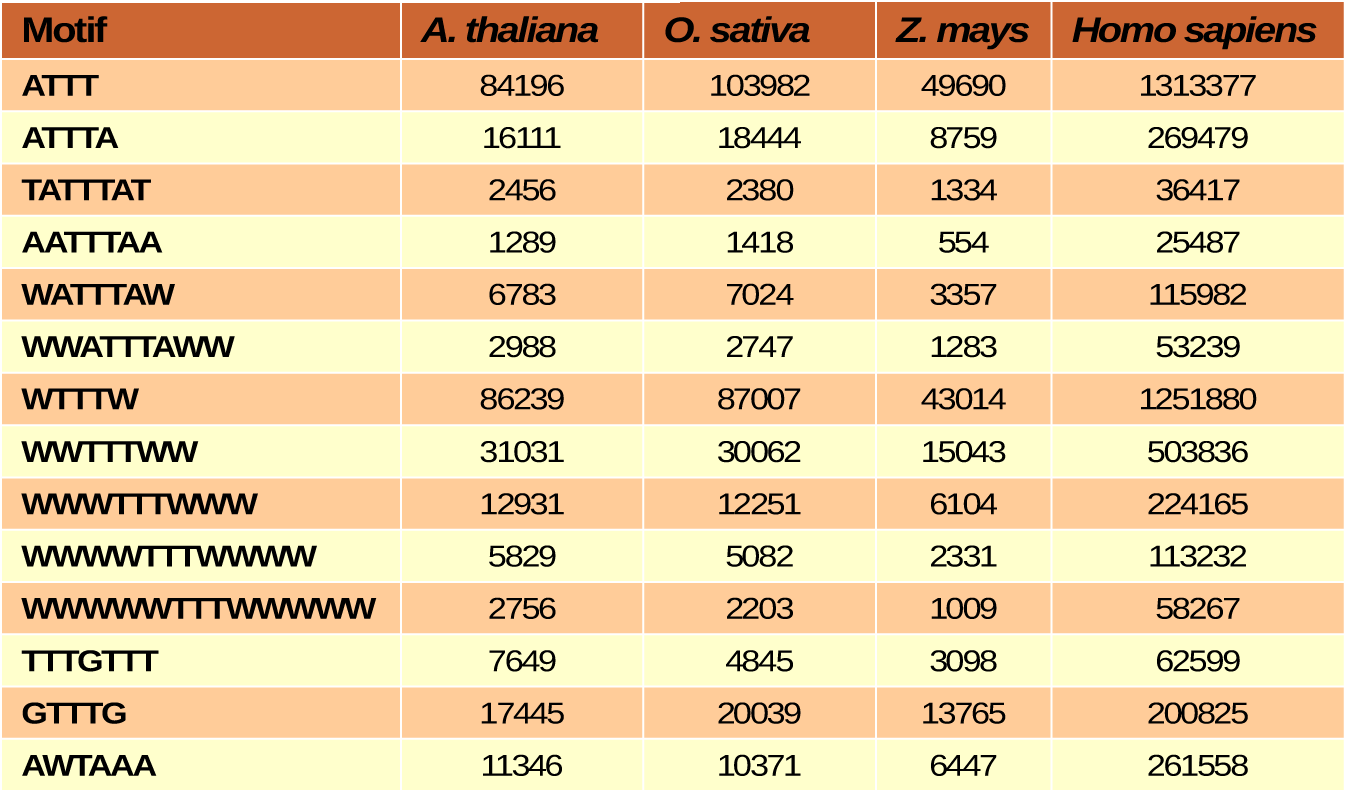
Number of ARE motifs present in 3’UTR of respective plant species and H.sapiens.

The magnitudes of genomes, transcriptome and 3’UTR of all the three plant species were disproportionate and incomparable with *Homo sapiens*. For the purpose of accessing the general propensity of ARE motifs in all species at the genome, transcriptome and 3’UTR level we standardized the size of the respective genome, transcriptome and 3’UTR by converting them to 100 MB (one unit) (Figure 3). Following size standardization, ARE motifs at a different stages can be compared realistically.

**Figure 3:**
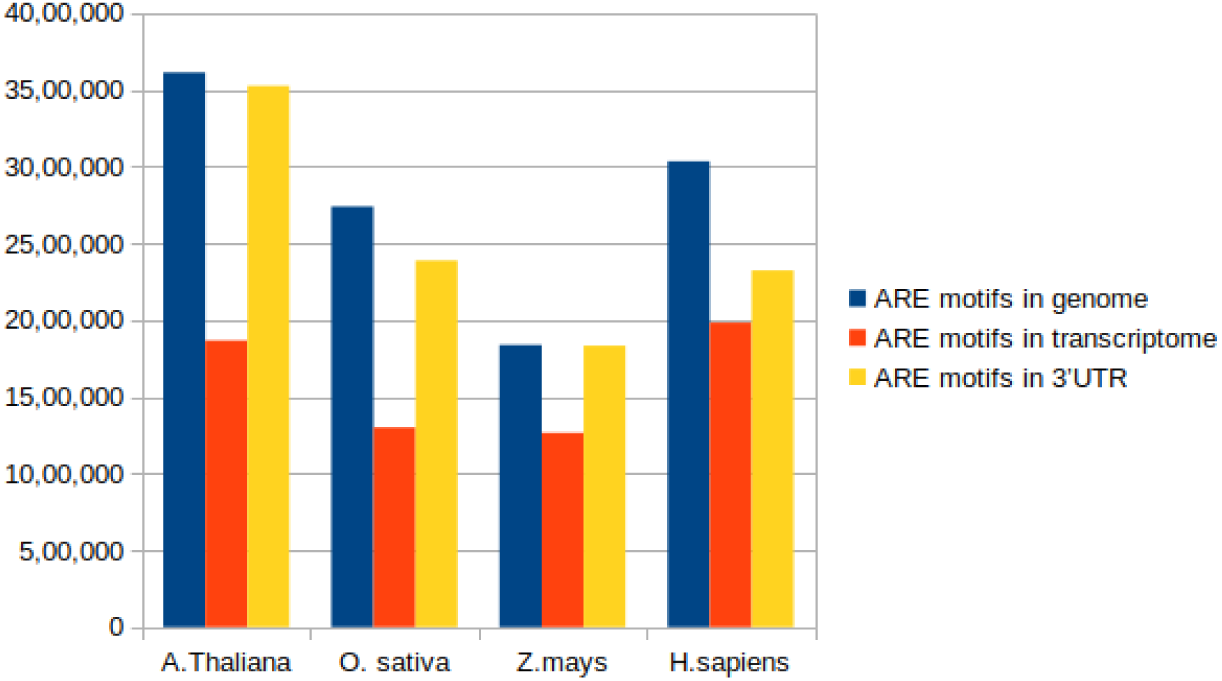
General trend of total ARE motifs at genome, transcriptome and 3’UTR level. Size at each level is taken as 100 MB (1 Unit) in order to make the data of ARE motifs comparable.

The present study indicated differential presence of ARE motifs in different plant species. But strong presence of ARE motifs at each level in all species under study cannot be denied. The most important inference deduced is that, similar to *H.sapiens*, plants also have more number of ARE motifs at 3’UTR than in per unit transcriptome. This is a breakthrough observation, which indicates the high probability of occurrence of ARE motifs in plants. The comparable magnitude of various ARE motifs at the genome, transcriptome and 3’UTR level in plants with *H. sapiens* indicates their possible roles in plant cell functioning similar to *H.sapiens*.

Additionally *in-vitro, in-vivo* and computational interventions are needed to acertain if the plants also have significant role of AU rich elements similar to mammals. The preliminary study conducted here categorically indicated the wide swath presence of AU rich elements in plants. Most prominent ARE motif and their role ascertained through further experimentation will open new avenues in the field of plant immunity and plant cell metabolism studies.

## Conclusion and future Prospects

The prevailing and significant ARE motifs are predicted computationally in three plant species and statistically proved to have a explicit presence in different plants species in the present genome-wide study. This is the first-ever study encompassing three different plants, showing the presence of AU rich elements at different genomic and transcriptomic levels. Existence of AU rich elements in plants could be used to study their role in plant immunity, stress tolerance, reproduction and productivity. The present study qualified to be an initiation point for further computational, statistical, in-vitro and in-vivo analysis.

Figure 4 provided a concrete pipeline for comprehensive computational and statistical extrapolation. This pipeline suggests ARE cluster analysis for enhanced interpretation of ARE motif occurrence and functioning. Understanding of localization and classification of ARE motifs is suggested to circumscribe 5’UTR and introns besides regions studied in the present case.

**Figure 4:**
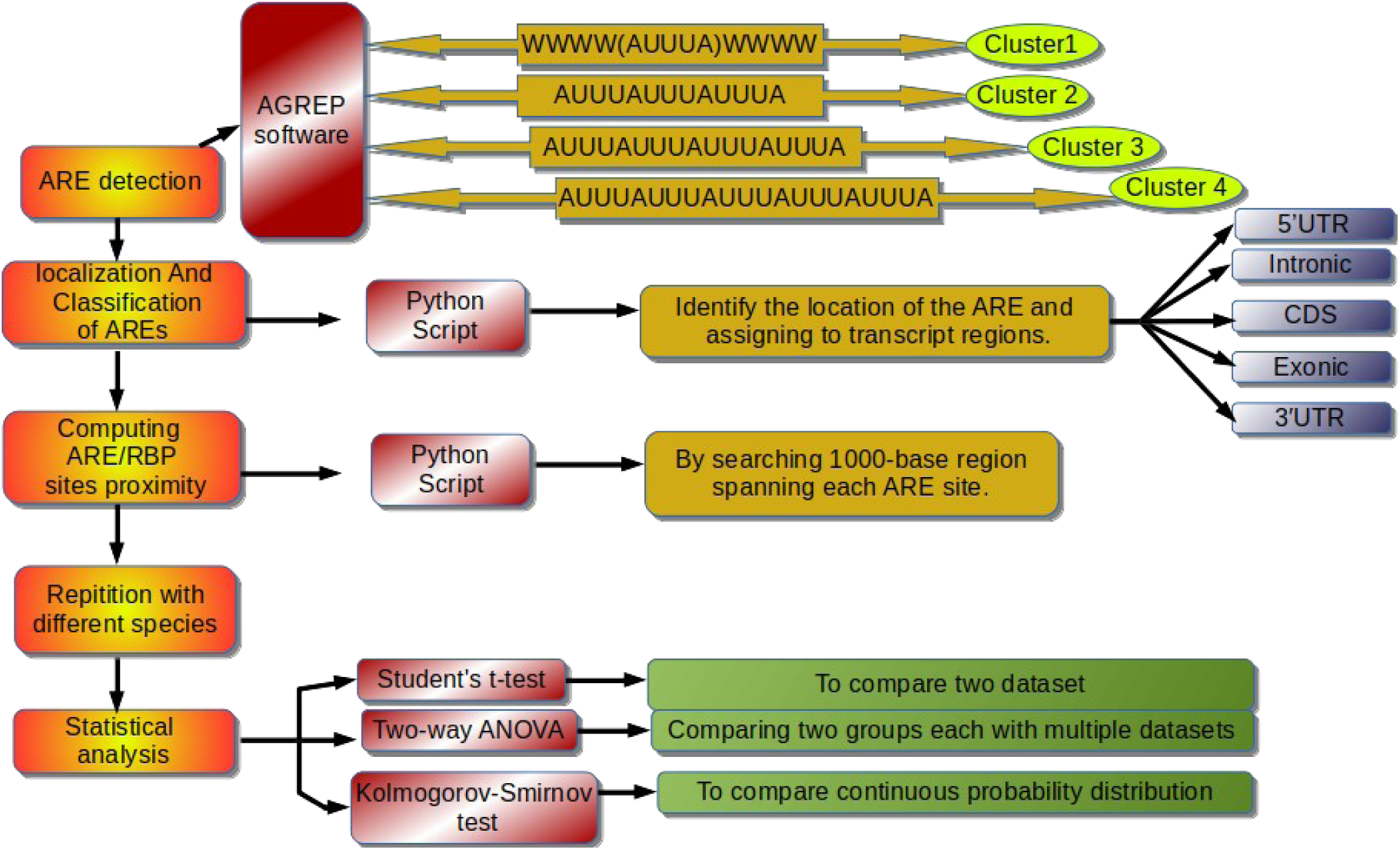
Pipeline of proposed computational analysis of genome-wide ARE motifs

ARE sites proximity might also be studied to get a extensive insight about the functioning of AU rich element directed mRNA decay. Understanding of appropriate proximity for functional ARE motifs will be of immense interest because cellular machinery needs space to get loaded on ARE signals. Student’s t-test, Two-way ANOVA test and Kolmogorov-Smimov test might be used for statistical analysis of elaborated results.

To get a comprehensible view regarding presence and significance of AU rich elements in plants study might be extended to encompass more number of plant species having complete genomes available and plants of immense economic importance e.g. *Triticum* aestivum, *Sorghum bicolour, Vitis vinifera, Medicago truncatula, Solanum tuberosum*.

